# Evaluation of melanocyte stem cell population in vitiliginous skin pre- and post-narrow-band UVB phototherapy

**DOI:** 10.1101/2024.02.06.579209

**Authors:** Abhik Dutta, Dyuti Saha, Manish Poojary, T S Nagesh, Colin Jamora

## Abstract

Vitiligo vulgaris is an autoimmune disorder characterized by the destruction of epidermal melanocytes leading to white lesions devoid of pigmentation. The destruction of epidermal melanocytes. Is driven by tissue-resident and circulatory immune cells that maintain an inflammatory milieu. Two percent of the global population suffer from vitiligo vulgaris, of which, prevalence is higher in the Indian population. Narrow-band UV-B (NBUVB) phototherapy is commonly used as an economical, non-invasive, and well-tolerated treatment modality. NBUVB promotes the migration of melanocyte stem cells (MelSC) to the epidermis, thus regaining pigmentation in affected individuals.

Despite the widespread use of NBUVB in secondary and tertiary clinics in India, the status of melanocyte stem cells and immune cells, pre-and post-NBUVB is not well understood. We observe that following NBUVB phototherapy, melanocyte stem cells (pax3+) re-populate the epidermis and express melanogenesis markers (Tyrosinase, Trp2, kit) concomitant with a decrease in CD3+ T-cells within the skin. Our study reaffirms the therapeutic efficacy of NBUVB in promoting re-pigmentation in an Indian cohort of vitiligo patients. Further, NBUVB is effective in reducing overall immune load, widely considered a major contributor to the pathogenesis of the disease.

## Introduction

Vitiligo Vulgaris is characterized by the complete loss of melanocytes and pigmentation from the epidermis, thereby leading to patches of de-pigmented skin. About 1-2% of the world population suffers from vitiligo and prevalence is as high as 4% within the Indian population ^1,2^. Patients suffering from vitiligo have a profound effect on their quality of life facing severe psychological and social problems thereby affecting their emotional and mental well-being.

The etiology of vitiligo remains complex and not fully understood. Accumulating evidence indicates the presence of stressed melanocytes in vitiligo patients, although the chemical or environmental triggers remain elusive. Stressed melanocytes overproduce reactive oxygen species (ROS), a by-product of melanin synthesis, which leads to an increase in cellular stress ^3^. Further, stressed melanocytes lead to the generation of damage-associated molecular patterns (DAMPs) that can induce inflammation through pattern recognition receptors (such as toll-like receptors) present on the surface of innate immune cells ^3^. Ultimately, the generation of an inflammatory milieu leads to the recruitment of melanocyte-specific CD8+ T-cells that are primarily responsible for the destruction of epidermal melanocytes leading to the characteristic depigmented lesions.

The treatment for vitiligo relies on replenishing the pool of epidermal melanocytes that are destroyed by cytotoxic CD8+ T-cells. Several treatment regimens are utilized to achieve this feat, these include – phototherapy-based approaches (UV-B-based), anti-inflammatory approaches (through corticosteroids and JAK-inhibitors), surgical approaches (hair grafting and micro-needling), or their combination. Among these, one of the frequently used, economical, well-tolerated, effective, and clinically approved methods for treating vitiligo is by exposing the depigmented region to narrow-band UV-B radiation (NBUVB) ^4,5^. NBUVB treatment is widely used in clinics and is often a central component of combinatorial therapy involving corticosteroids, JAK inhibitors, and psoralen among others ^6^.

Despite its widespread use in clinics, a thorough investigation into the melanocyte behavior in the re-pigmentation process following NBUVB treatment is currently lacking among the Indian population. In this study, we investigate the changes in melanogenesis, melanocyte stem cell (MelSC) population, and immune profile in Indian vitiligo patients treated with NBUVB.

## Results

NBUVB therapy was performed on adult Indian patients with stable Vitiligo vulgaris according to previously described methods ^5,7^ at a tertiary care center with Institutional Ethics Committee approval. Patient demographics including age, duration of vitiligo, and location is summarized in Table 1. Punch biopsies (3mm) were obtained from vitiligo patients before starting the NBUVB treatment regimen and 3 months post-treatment. A biopsy from the contralateral to the affected region was taken from each patient to serve as a baseline (Figure 1A). Following 3-months of NBUVB treatment alone, perifollicular re-pigmentation was observed in patients.

**Table 1.** Clinical characteristics of Vitiligo patients.

| Patient No. | Age | Sex | Body site involvement | Vitiligo disease duration (in years) | Duration of narrow-band UVB treatment (in months) | Previous treatments, if any |
| --- | --- | --- | --- | --- | --- | --- |
| 1 | 24 | M | chest, back and limbs | 4 | 3 months | topical medications |
| 2 | 30 | M | nape of neck | 2 | 3 months | nil |
| 3 | 50 | M | lower abdomen | 2 | 3 months | topical medications |
| 4 | 52 | M | face, dorsum of hands | 5 | 3 months | topical medications |
| 5 | 22 | M | left side of chin | 3 | 3 months | topical medications |
| 6 | 50 | F | upper limbs and trunk | 10 | 3 months | topical medications, ayurvedic and homeopathic medicines |
| 7 | 40 | F | back | 1 | 1 month | topical medication |
| 8 | 29 | M | upper and lower limbs | 3 | 1 month | topical medication |
| 9 | 30 | M | generalised vitiligo | 5 | 3 months | systemic steroids, topical medications |
| 10 | 22 | F | upper and lower limbs | 2 | 2 months | topical medication |
| 11 | 62 | M | face, dorsum of hands, feet | 1 | 1 month | topical medication |
| 12 | 25 | F | lower abdomen | 10 | 2 months | topical medication |
| 13 | 39 | F | legs | 3 | 2 months | topical medications |

**Figure 1.**
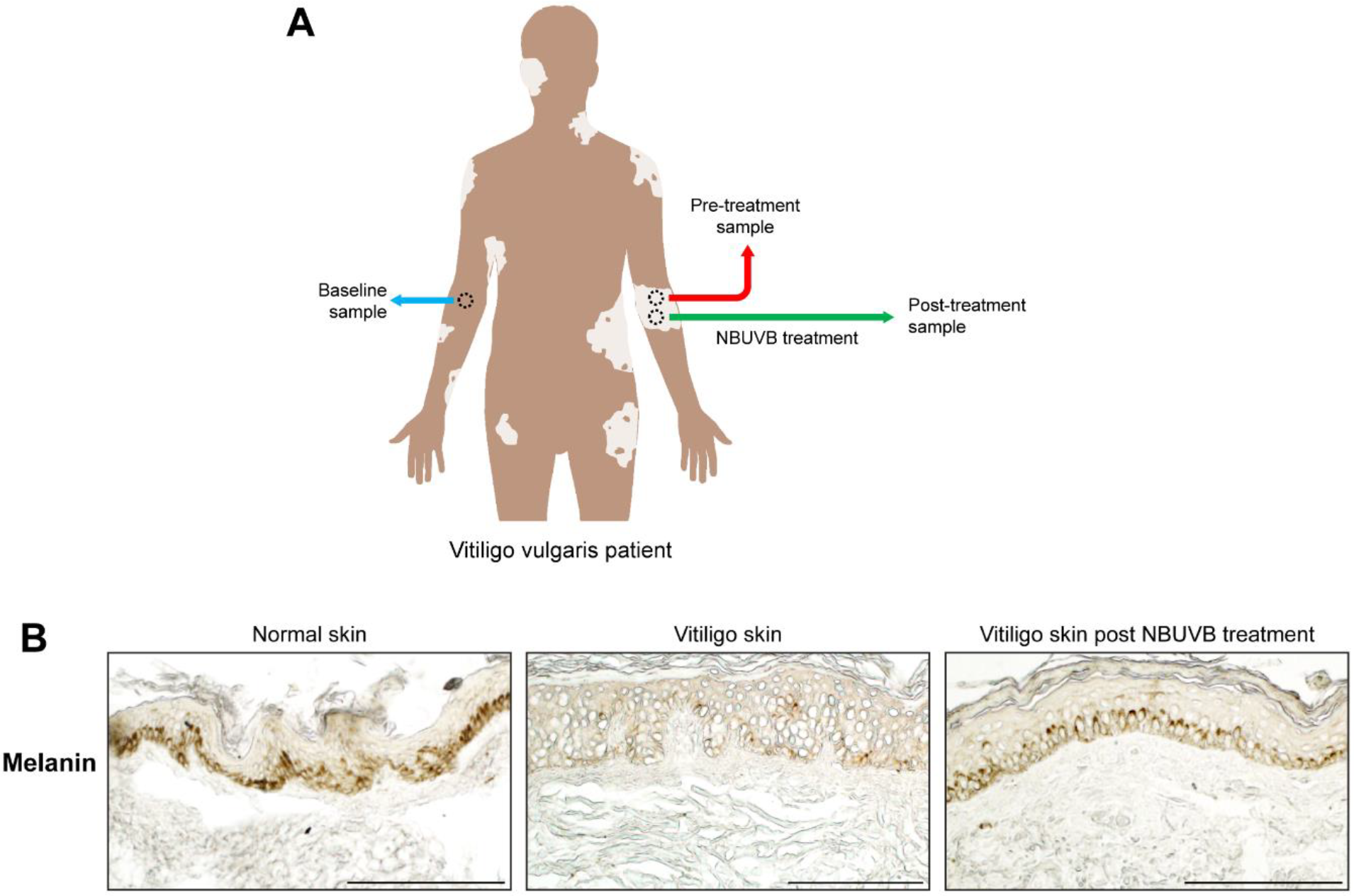
NBUVB treatment promote re-pigmentation in adult Indian vitiligo patients. (A) Schematic of skin biopsy samples obtained from vitiligo vulgaris patients. Three punch biopsies were obtained from each patient – contralateral biopsy (blue), pre-treatment biopsy (red) and post-treatment biopsy (green) (B) Bright field images of melanin deposition in skin sections of normal, vitiligo skin and vitiligo skin post-NBUVB treatment. Vitiligo patients treated with NBUVB (right panel) restores the pigmentation significantly, comparable to that of normal skin. Images are representative from n=5 patients. Scale bar = 100μm.

### NBUVB treatment causes re-appearance of melanocytes in the epidermis and restores pigmentation in Indian vitiligo patients

NBUVB treatment has been approved as a tertiary level of treatment for vitiligo vulgaris by the Indian Council of Medical Research. To investigate the status of pigmentation following NBUVB treatment, we performed histological analysis of skin sections from vitiligo patients pre- and post-NBUVB treatment. This analysis revealed that melanin pigment was deposited in the basal layer of the epidermis post-NBUVB treatment (Figure 1B).

Since NBUVB treatment alone is sufficient to promote re-pigmentation of the affected region, we then sought to assess the gene expression profiles of genes involved in melanogenesis. Several transcription factors and intracellular proteins serve as critical checkpoints during melanogenesis. Notable among these are – KIT (which promotes proliferation and migration of melanocytes), TYR (which encodes an enzyme Tyrosinase which catalyzes dihydroxyphenylalanine in the melanin synthesis pathway), and TRP-2/DCT (which encodes the enzyme dopachrome tautomerase involved in modification of pigment color) ^8–10^. Melanogenesis is perturbed in vitiligo patients and the activity of the aforementioned genes is drastically reduced in vitiligo patients ^11,12^. We performed quantitative gene expression analysis from skin biopsies obtained from vitiligo patients pre- and post-NBUVB. A contralateral skin biopsy from an unaffected region (having normal pigmentation) served as a baseline for each patient. Quantitative gene expression analysis revealed that the mRNA levels of KIT, TYR, and TRP-2/DCT were significantly upregulated in NBUVB post-treated vitiligo skin compared to pre-treated vitiligo skin (Figure 2B).

**Figure 2.**
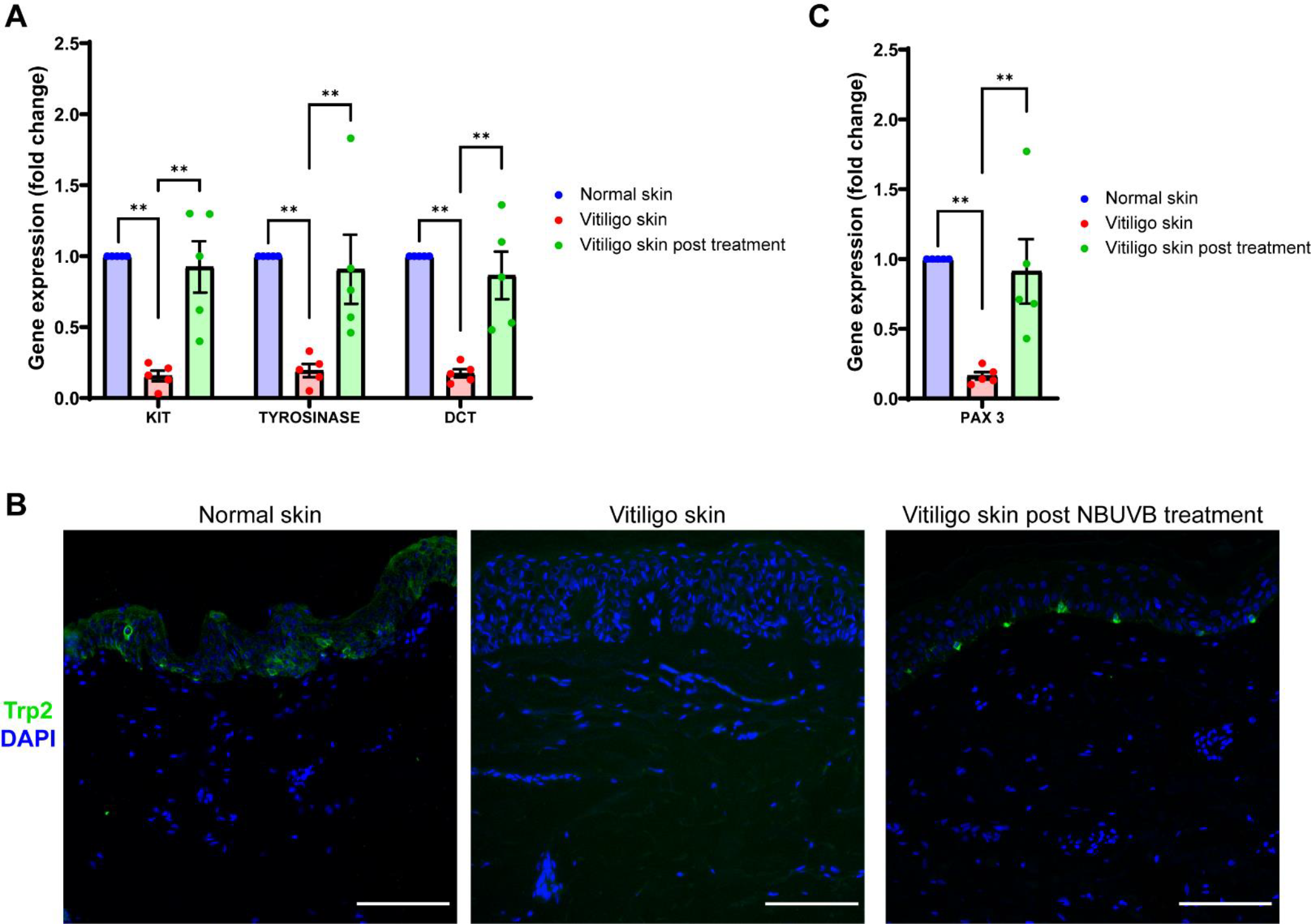
NBUVB treatment promote re-appearance of melanocytes and melanogenesis in vitiligo patients. (A) Quantitative gene expression analysis for melanogenesis genes –Kit, Tyrosinase and Dct. Gene expression is normalized to normal skin levels and represented as fold change. Statistical analysis was done using Mann-Whitney U Test (^**^p < 0.01; n=5 patients). (B) Immunofluorescence staining of pan melanocyte marker – Trp2 (in green). In normal human skin (left panel), the melanocytes reside in the epidermal compartment, which are entirely absent in vitiligo skin (middle panel) and re-appear in vitiligo patients treated with NBUVB (right panel). Images are representative from n=5 patients. Scale bar = 100μm. (C). Quantitative gene expression analysis for melanocyte stem cell gene – pax3. Gene expression is normalized to normal skin levels and represented as fold change. Statistical analysis was done using Mann-Whitney U Test (^**^p < 0.01; n=5 patients).

Pigment-producing melanocytes arise from MelSC that reside in the hair bulge region of the hair follicle ^13^. Evidence from mouse and human clinical data suggests that melanocyte stem cells (MelSC) migrate to the epidermis in response to NBUVB and differentiate into melanin-producing melanocytes ^14,15^. We next assessed the status of MelSC using a pan melanocyte marker – Trp2, expressed by MelSC and differentiated melanocytes, and gene expression of Pax3 (a transcription factor that regulates the proliferation and differentiation of melanocyte stem cells). We observe that NBUVB is capable of restoring the melanocytes in the epidermis that are completely absent in skin sections of vitiligo patients (Figure 2B). Furthermore, a concomitant increase in gene expression of Pax3 following NBUVB therapy, confirms that MelSC are required for re-pigmentation of vitiligo patients (Figure 2C).

Altogether, our analysis suggests that MelSC re-populate the epidermis and differentiates to produce melanin pigment in Indian vitiligo patients treated with NBUVB.

### Vitiligo patients post-NBUVB treatment exhibit decreased immune load

Melanocyte-specific cytotoxic CD8+ T-cells proximal to the epidermal-dermal region are responsible for the destruction and clearance of epidermal melanocytes ^16,17^. Consequentially, the border of depigmented lesions exhibits increased T-cell infiltration and increased pro-inflammatory cytokine and chemokine levels – IFNγ, TNFα, IL-17, CXCL9 and CXCL10 ^16^.

To investigate the status of T-cells following phototherapy, we performed immunostaining on skin sections of vitiligo patients pre- and post-NBUVB. In accordance with existing reports ^16,17^, we observe a significant increase in total number of CD3+ T-cells in de-pigmented region of vitiligo patients compared to contralateral region from the same patients (Figure 3A). Following NBVUB treatment, we observe a significant reduction in CD3+ T cells similar to the levels of contralateral region (Figure 3B).

**Figure 3.**
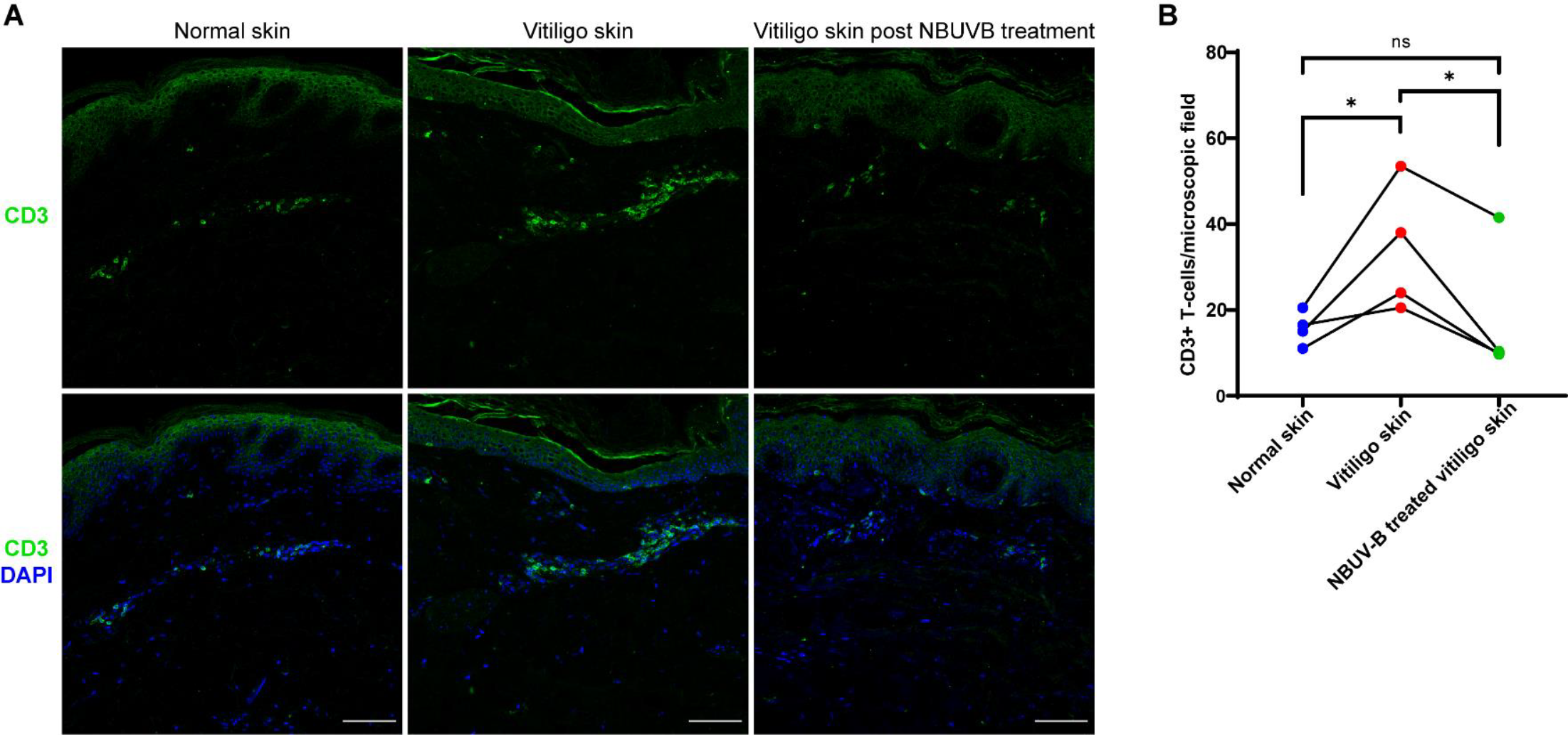
Vitiligo patients post-NBUVB treatment exhibit decreased immune load. (A) Skin sections from biopsies obtained from vitiligo patients from contralateral region (normal skin), pre-treated (vitiligo skin) and post-NBUVB treatment were immunostained for pan T-cell marker – CD3 (in green) and DAPI (in blue). Skin biopsies obtained from post-NBUVB treatment show a drastic decrease in CD3 T-cells compared to pre-treated vitiligo skin sections indicating immunosuppressive effect of NBUVB. CD3+ T-cells per microscopic field is quantified in (B). Images are representative from n=4 patients. Scale bar = 100μm. Statistical analysis was done using Mann-Whitney U Test (^*^p < 0.05; n=4 patients).

Taken together, we observe that NBUVB treatment is sufficient to induce re-pigmentation and decrease the overall immune load in Indian vitiligo patients.

## Discussion

Existing studies involving the Indian vitiligo population are largely prospective studies with only limited information available regarding the status of melanocytes, thus, necessitating a comprehensive evaluation of vitiligo patients post-NBUVB phototherapy. In our study, we report that NBUVB phototherapy alone is sufficient to induce re-pigmentation of stable generalized vitiligo cases among the Indian population. Vitiligo patients treated with NBUVB have increased expression of MelSC genes and re-appearance of epidermal melanocytes. Further, the melanogenesis pathway is upregulated post-phototherapy ultimately leading to deposition of melanin in the epidermal compartment. Concomitant to the re-pigmentation of vitiligo skin, we observe a reduction in skin-infiltrating T-cells. Taken together, our study sheds light on the mechanism of action of NBUVB phototherapy and reaffirms the efficiency of NBUVB phototherapy for Vitiligo vulgaris as reported by previous studies ^5,18–20^.

The autoimmune destruction of epidermal melanocytes serves as an attractive target for remediating the progression of vitiligo. Topical corticosteroids or calcineurin inhibitors are administered to minimize the spread of active lesions and stabilize the condition ^21^. In cases of rapidly progressing lesions, systemic reduction of inflammation using oral corticosteroids (such as dexamethasone) is administered to the patient, however, adverse metabolic side effects have been reported from long-term use ^22^. Emerging approaches target the IFNγ-CXCL10 chemokine axis by interfering with the JAK-STAT signaling pathway. Oral administration of tofacitinib (pan-JAK inhibitor) and ruxolitinib (JAK 1/2 inhibitor) promote only temporary re-pigmentation in vitiligo patients ^23^. Despite the mounting evidence of immune-mediated destruction of melanocytes in vitiligo pathogenesis, the reduction of inflammation alone is not sufficient to induce repigmentation. However, inhibition of the overall immune burden can fasten the rate of repigmentation when used in combination with NBUVB phototherapy ^6^. These results underscore the value and importance of NBUVB phototherapy and hint towards hitherto unknown mechanisms of action of NBUVB.

MelSC are immature reservoirs of melanocytes that reside in the hair bulge region of the hair follicle^13^. Following NBUVB treatment of vitiligo patients, MelSC migrates along the hair follicle and emerges into the epidermis, a process termed perifollicular re-pigmentation^14,24^. Consequently, clinical studies have reported better outcomes (more re-pigmentation) upon NBUVB phototherapy in regions with a higher density of hair follicles (perifollicular repigmentation) as opposed to regions devoid or scarce hair follicles (marginal and diffuse repigmentation) ^25^. Thus, understanding the mechanism of action of NBUVB in promoting MelSC migration can fuel the development of novel therapeutics independent of NBUVB exposure.

## Materials and methods

### Human subjects

A total of 13 adult patients with vitiligo were included in the study. Written informed consent for the procedure was obtained from the participants. Six patients dropped out of the study. Out of the 7 patients who completed the NBUVB therapy for 3 months, biopsies were analysed from 5 patients. Demographic details of all patients, including age, duration of disease and the treatment history, are summarized in Table 1. Approval to extract patient samples with informed consent was obtained from the Ethical Committee of Sapthagiri Institute of Medical Sciences and Research Center (Bangalore, India). Analysis of patient skin samples in the Jamora lab (Bangalore, India) was approved by the Institutional Ethics Committee of the Institute for Stem Cell Science and Regenerative Medicine (Bangalore, India) (Human Ethics Approval Certificate number inStem/IEC-8/003).

### RNA isolation, cDNA synthesis, and qPCR

Single 3mm punch biopsies from control skin, Vitiligo skin before treatment and Vitiligo skin 3 months post NBUVB treatment from each recruited patient were collected in RNA stabilization buffer. Individual punch biopsies were homogenized in 1 ml RNAiso Plus reagent (DSS Takara Bio India Pvt. Ltd, Catalog number 9109), and RNA was isolated according to the manufacturer’s instructions. cDNA was prepared using PrimeScript cDNA Synthesis Kit (DSS Takara Bio India Pvt, Ltd; Catalog number 2680A) according to the manufacturer’s instructions. qPCR was performed using PowerUp SYBR Green Master Mix (Applied Biosystems, Thermo Fisher Scientific, Catalog number A25742) in CFX384 machine (Bio-Rad Laboratories). β-Actin expression was used for normalization. The primers used are listed in Table 2.

**Table 2.**
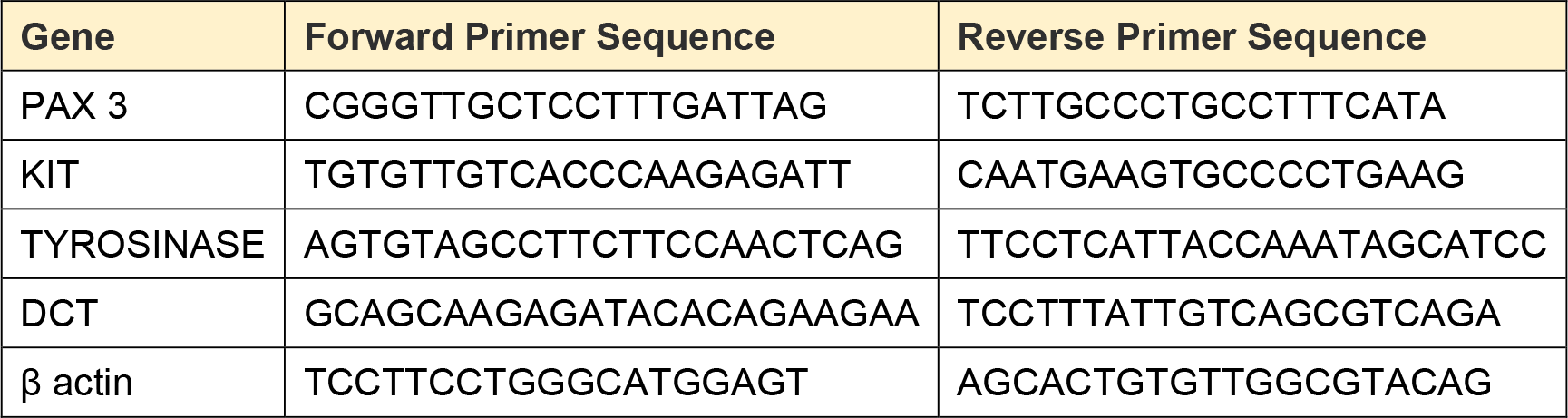
qPCR primer sequences.

### Histology and immunohistochemistry

A 3mm skin biopsy specimen was taken from control skin, vitiligo skin prior to treatment, and vitiligo skin post-NBUVB treatment. Skin biopsies were fixed in 10% neutral buffered formalin for 2 hours at room temperature. After sequential dehydration using increasing concentrations of ethanol (50%, 60%, 80%, 95%, and 100%) and xylene, the skin biopsies were embedded in paraffin wax. 8µm sections were used for bright field imaging and immunohistochemistry. Paraffin sections were deparaffinized and antigen unmasking was performed in antigen retrieval buffer (10mM sodium citrate buffer pH 6) in a water bath maintained at 65°C. Tissue sections were then blocked (5% normal donkey serum, 1% bovine serum albumin, 0.3% Triton X-100) for 1 hour at room temperature followed by overnight incubation at 4°C with primary antibody against TRP2/DCT (1:200, Abcam, catalog number ab74073). Alexa fluor-647 labeled secondary antibody (1:300, Invitrogen, catalog number A21244) was used to detect the primary antibody. DAPI was used to stain nuclei. Imaging was performed on an Olympus IX73 microscope or FV3000 confocal microscope and the images were analyzed on ImageJ (Fiji) software.

### Statistical analysis

Comparison of two groups was done using non-parametric Mann-Whitney U test. GraphPad Prism (8.4.2) was used to perform all statistical analysis. p-value < 0.05 was considered significant.

